# Bispecific Coiled-Coil Reporters Enable Rapid, Multiplexed Detection of Mycoplasma via Toehold Switch Sensors

**DOI:** 10.64898/2026.05.28.728408

**Authors:** Yudan Li, Nery R. Arevalos, Amit Eshed, Shannon Newell, Carlye Frisch, N. Rebecca Kang, Xiao Wang, David A. Brafman, Alexander A. Green

## Abstract

Cell-free synthetic biology offers a rapid prototyping environment for molecular diagnostics, yet the utility of these systems is often limited by the slow kinetics and restricted multiplexing capabilities of traditional protein reporters. Here, we report the BINOCULAR (BIspecific Novel Orthogonal Coiled-coil-Using LAteral flow Reporter) system based on supramolecular coiled-coil peptide conjugates that dramatically accelerates signal generation and enables robust multiplexing in cell-free transcription-translation reactions. Using BINOCULARs as toehold switch outputs, we achieve visible readout times of less than 5 minutes in lateral flow assays, achieving a ≥12-fold increase in speed compared to standard cell-free reporters like β-galactosidase or fluorescent proteins. To demonstrate the utility of this platform, we integrated the bispecific reporters with loop-mediated isothermal amplification (LAMP) and a lateral flow assay for the detection of *mycoplasma*, a common adventitious agent in biomanufacturing. The integrated system demonstrates exceptional sensitivity, achieving a detection limit of 1 copy/µL for *Mycoplasma* 16S rRNA and a mammalian internal control gene. Cell-free assay results are available within 5 minutes following a 20-minute amplification step, and we successfully achieved the simultaneous detection of three distinct targets in a single reaction. Crucially, the platform maintains reliable performance in complex matrices and after lyophilization, making it more accessible for point-of-use need. Our findings suggest that the modular BINOCULAR architecture can overcome current bottlenecks in cell-free sensing systems, providing a versatile and scalable framework for rapid, point-of-use screening across clinical and industrial applications.

## INTRODUCTION

Cell-free synthetic biology has emerged as a transformative framework for molecular diagnostics, offering a rapid, adaptable environment for prototyping and deploying genetic circuits^1–3^. By harnessing the transcriptional and translational machinery of the cell in an in vitro format, researchers have created complex biosensors that operate independently of living hosts, significantly increased the assay accessibility, and facilitated the use of non-canonical components. A prominent implementation of this field is the integration of cell-free transcription-translation (TX-TL) systems with programmable RNA sensors, such as toehold switches.^4^ In previous work, we developed a set of alternative nucleic acid detection strategies leveraging cell-free transcription-translation (TX-TL) reactions. By lyophilizing cell-free components onto paper substrates, we preserved their biological activity, allowing for simple reactivation via water reconstitution. By coupling these components with toehold switch sensors encoding full-length or split ß-galactosidase (lacZ), we demonstrated the rapid, low-cost, and sensitive detection of pathogens such as Zika virus and norovirus^5–8^. This approach significantly reduced the equipment and reagents required for amplicon readout, facilitating convenient testing both inside and outside laboratory settings. Moreover, it enabled the recognition of pathogens and drug-resistance mutations with enhanced performance using mutation-specific sensors^9^ and molecular logic systems^10^,and facilitated assay readout with novel output schemes, including electrochemical systems^11^, home glucose monitors^12^, and wireless sensors^13^.

Despite these advances, the utility of cell-free diagnostics is frequently constrained by the limitations of traditional reporter systems. Standard output modalities typically rely on the expression of large enzymes, such as LacZ, or fluorescent proteins, such as GFP. While reliable, these reporters possess inherent disadvantages: they are relatively large proteins requiring significant TX-TL time for folding and maturation, often resulting in signal-to-readout delays of 30 to 120 minutes^6,7^. Furthermore, substrate spectral overlap has severely hindered colorimetric outputs’ potential for multiplexing^14^, while fluorescence-based methods require specialized plate readers or portable optics, limiting their use in true point-of-need settings. Although alternative schemes involving electrochemical sensors or glucose monitors have been developed to improve sensitivity and multiplexing, they often require additional, specialized instrumentation, complex sample preparation, and longer readout time^12^.

Therefore, there is a critical need to reduce readout times and simplify detection formats to make cell-free diagnostics commercially and clinically viable. To address this, we hypothesized that the development of a modular, bispecific reporter system leveraging the rapid expression kinetics of small peptides rather than larger proteins could dramatically enhance the speed and scalability of cell-free assays. Supramolecular coiled-coils represent a particularly promising tool for this purpose^15,16^. These minuscule peptides (typically <50 amino acids) can be expressed in extremely short timeframes and form stable, predictable multimeric bundles via specific heptad repeats. Indeed, our lab has previously exploited cell-free expression of coiled-coil-nanobody conjugates for rapid detection of different protein antigens^17^. Importantly, the existence of libraries of orthogonal coiled-coil pairs^18^ offers a direct and scalable tool for a “plug-and-play” architecture that can facilitate simultaneous detection of multiple targets without signal interference, which was previously unachievable.

The practical utility of such an accelerated, multiplexed platform is well suited to address the demands of modern biomanufacturing^19^, where the integrity of complex bioprocesses is under constant threat from adventitious agents^20,21^. For example, the manufacturing of vaccines and monoclonal antibodies is inherently vulnerable to biological contaminants that can bypass traditional safeguards. Among these threats, mycoplasma is notoriously problematic. Their small size (0.15–0.3 µm) and lack of a cell wall allow them to bypass standard 0.22 µm sterilization filters, while their cryptic nature allows them to proliferate to high titers without causing visible turbidity^22–24^.

Historically, the biopharmaceutical industry has relied on “gold-standard” culture-based assays for mycoplasma detection^25^. These methods are established by global pharmacopoeias for maximum sensitivity to ensure no viable contaminant is missed, but can require up to 28 days for a final result^25,26^. While modern molecular methods like qPCR have reduced this timeline to several hours, they remain expensive, laboratory-bound, and susceptible to matrix interference from complex cell culture media^27,28^. Although isothermal methods like LAMP have been explored to overcome the need for lengthy thermal cycling, these assays still rely on fluorescence readers or gel electrophoresis as specificity filters^29,30^. Thus, there is a clear need for rapid, low-cost, and easy-to-deploy mycoplasma detection platforms, particularly those that support real-time quality assurance and reduce dependency on centralized testing infrastructure.

Here, we report the development of BIspecific Novel Orthogonal Coiled-coil-Using LAteral flow Reporters (BINOCULARs) that bridge the gap between high-sensitivity nucleic acid detection and rapid visual readout for cell-free reactions (Figure 1). By employing a bispecific protein reporter composed of a dual coiled-coil fusion peptide, we integrated BINOCULARs with lateral flow strips to achieve an instrument-free, rapid visual readout for cell-free reactions. Our platform achieves a sensing-to-readout time of 5 minutes, which is over 12-fold faster than standard cell-free reporters, while maintaining qPCR-level sensitivity for mycoplasma and other adventitious agents. Furthermore, by leveraging orthogonal coiled-coil pairs and their corresponding capture probes, we demonstrate the simultaneous detection of three distinct targets in a single reaction. Finally, we show that the platform exhibits significant robustness; it remains sensitive in diverse sample matrices and retains activity after lyophilization and storage for 35 days. This modular architecture not only provides a solution for bioprocess safety but also offers a versatile framework for cell-free sensing across a diverse range of applications.

**Figure 1.**
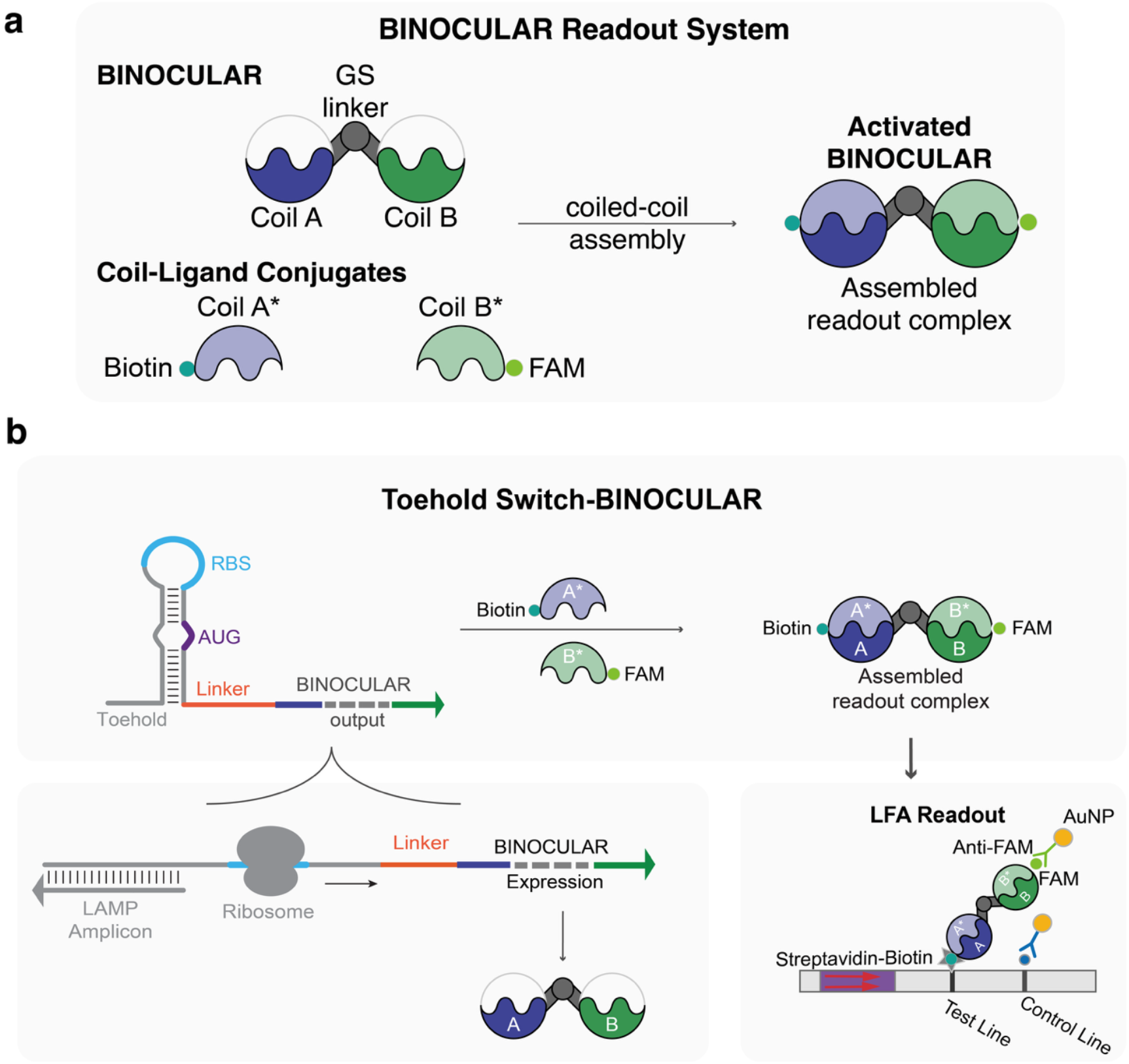
Schematic illustration of the BINOCULAR system and the Toehold Switch-BINOCULAR assay for fast visible nucleic acid detection. (a) BINOCULARs consist of two orthogonal coil domains that flank a central glycine-serine (GS) linker and enable binding to a pair of cognate coils for lateral flow readout. (b) A toehold switch sensor encoding the BINOCULAR output is added to a cell-free reaction. Upon detection of a target by the toehold switch, expression of the BINOCULAR having coils A and B is activated. Following cell-free incubation, cognate binding partners Coil A* (conjugated to a capture probe, e.g., biotin) and Coil B* (conjugated to a reporter, e.g., FAM) are supplied to the reaction. The resulting product is applied to a lateral flow strip, where captured BINOCULAR/coiled-coil complexes accumulate at the test line to generate a visible signal.

## RESULTS

### Toehold Switch Design, Screening, and Selection

Based on previous literature^31^, we first identified a highly conserved 16S rRNA region specific to *Mycoplasma* that covers 90% of all known species and is suitable for isothermal amplification and toehold switch-based detection. *Mycoplasma* genome sequences were obtained from the NCBI Bacterial 16S Ribosomal RNA RefSeq Targeted Loci Project (Accession PRJNA33175) database. The target region was further refined to a 519-nucleotide sequence for subsequent sensor design. A similar process was performed to identify a universally expressed mammalian cell target sequence, the *PUM2* gene, to serve as an internal control for cell lines widely used in biomanufacturing. All target sequences are included in Supplementary Table 1.

Toehold switches for detection of the target sequence were then generated based on the NUPACK-based second-generation toehold switch algorithm^4,32,33^. We designed a library of 96 toehold switches (48 for each target), initially utilizing full-length LacZ as the reporter output for functional sensor screening. All tested toehold switch sensor sequences are included in Supplementary Table 2. All DNA encoding the toehold switches was cloned into vectors (Addgene #11882) upstream of the lacZ open reading frame. After sequence confirmation, sensor plasmids were screened in 5-µL liquid-phase cell-free reactions supplemented with a lacZ chromogenic substrate chlorophenol red-β-D-galactopyranoside (CPRG). The cleavage of the chromogenic substrate was monitored by optical density (OD) at 570 nm using a plate reader.

The toehold switch ON/OFF ratio was calculated based on the difference in OD570 between samples with the cognate trigger (ON) and those with a non-cognate trigger (OFF) (Figure 2a, Supplementary Figure 1). Sensors exhibiting an ON/OFF ratio >2 were shortlisted for verification, and the three top-performing sensors for each target were selected for further optimization.

**Figure 2.**
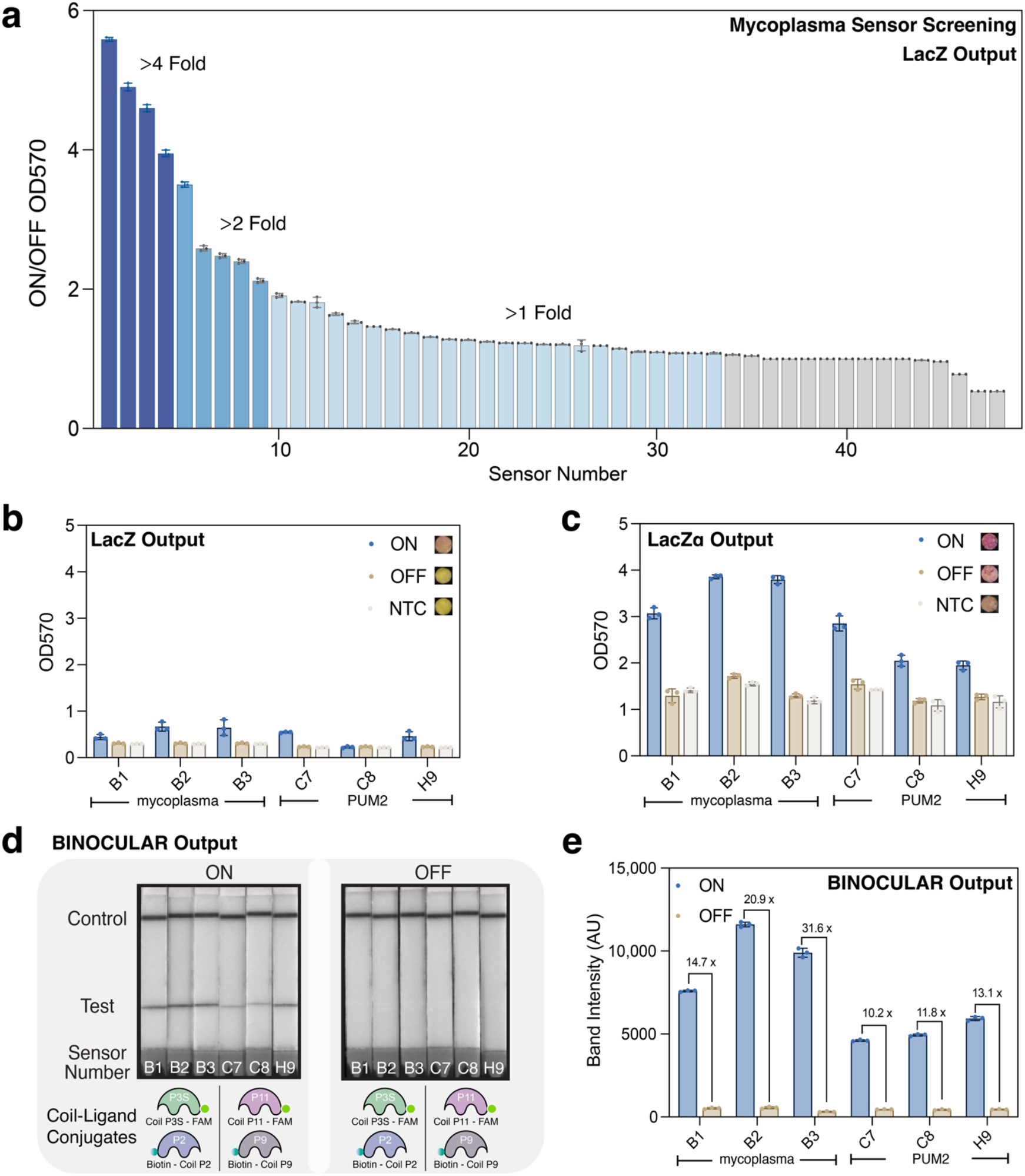
Screening of mycoplasma and PUM2 toehold switches with different reporter proteins. (a) Screening of functional toehold switch sensors with LacZ output. OD570 fold change obtained 2 hours of cell-free reaction for 48 toehold switch sensors determined in the presence or absence of 5 µM of cognate mycoplasma 16S ribosomal RNA gene target. The signal fold change was calculated by dividing the relative OD570 of the ON state by the OFF state. (b, c) OD570 fold change of 3 top-performing sensors at 30 minutes of the reaction for each target with full-length LacZ output (b) or LacZα output (c), with representative paper-based test color changes for sensor B2 on the right. B1, B2, B3 are mycoplasma sensors; C7, C8, H9 are PUM2 control sensors. (d) Representative image of one set of lateral flow strips analyzed in (e), which were used to quantify band intensities for the performance of coiled-coil output sensors. Sensor numbers and corresponding coil-ligand conjugates for each sensor are listed at the bottom of the frame. (e) Performance of the same sensors sequences in (d) with coiled-coil output after 5 minutes of cell-free incubation against a cognate target. Final concentration of target is 5 µM for all experimental groups. Error bars indicate standard deviation across n = 3 independent technical replicates.

### Development of a Rapid, Leakage-Free Reporter for Instrument-Free Readout

Although the top-performing sensors achieved saturation with low leakage within 2 hours, we sought to further accelerate the assay. Previously, our group reported that sensors utilizing the LacZ α-peptide (LacZα) reporter could achieve a readout in 33 minutes, significantly faster than the 56 minutes required for full-length LacZ^7^. This approach leverages the self-assembling nature of the enzyme: instead of expressing the entire ∼2.7 kb full-length gene, the system expresses a short 174 bp LacZα subunit, while the complementary LacZω peptide comprising the remaining ∼970 residues is supplied in the reaction. This strategy drastically reduces the transcription and translation time required.

Under identical reaction conditions at the 30-minute mark, sensors expressing LacZα exhibited a much greater signal change compared to those utilizing full-length LacZ (Figure 2b-c). However, this homologous reporter architecture presented a leakage issue, resulting in a higher background signal in the absence of the target for LacZα. This background signal was likely caused by low levels of reporter produced in the sensor’s OFF state. The enzymatic nature of the LacZ output dramatically amplifies these trace amounts of basal expression, as each translated enzyme molecule converts numerous substrate molecules. This type of reporter makes the system highly sensitive to even the slightest sensor leakiness.

To overcome the leakage inherent in the split-enzyme system and achieve a standardized, rapid readout, we sought to develop an easy-to-translate reporter that could be integrated with a lateral flow assay (LFA) detection system. Lateral flow assays employ two critical components: (i) mobile analyte recognition elements, which bind the target molecule; and (ii) stationary capture elements, which are immobilized on the strip to generate the readout signal. Typically in universal lateral flow strips, these capture systems utilize chemical moieties (e.g., biotin, FAM) rather than proteins for more versatile application^34^. These moieties facilitate the localization of the analyte-probe complex at the test line, causing the accumulation of reporter particles such as gold nanoparticles (AuNPs) and the formation of a visible band. The necessity for using AuNPs and lateral flow assay arises from the inherent sensitivity limitations of direct protein reporters. While strong visual detection of fluorescent or chromogenic proteins generally requires micromolar concentrations, which is often difficult to reach within the abbreviated timeframes of cell-free reactions, AuNPs possess exceptionally high extinction coefficients, typically ranging from 10^8^ to 10^10^ M^-1^cm^-1^.^35,36^ This high optical density yields a clear, visible signal from even picomolar concentrations of captured analyte, offering a sensitivity advantage of several orders of magnitude over direct protein visualization^37^. However, these chemical moieties cannot be directly produced via biological translation, which makes them incompatible as direct outputs for a genetically encoded toehold switch sensor. This limitation necessitates a reporter design that is fully proteinaceous yet functions as a specific capture agent.

To address this challenge, we exploited pairs of coiled coils in BINOCULARs to enable readout on lateral flow strips. BINOCULARs were designed as fusion proteins composed of two orthogonal coiled-coil peptides tethered via a flexible glycine-serine linker, where each coil comprised one half of a coiled-coil pair (Figure 1). The resulting small fusion protein is encoded by a 243-bp long DNA, which is about an eleventh of the length of lacZ. Further, because of the bispecificity of BINOCULARs, the orthogonality of the coils, and absence of multi-turnover enzymatic activity, we expected this output would create less leakage in comparison to the split LacZ architecture while still providing excellent sensitivity due to the robust LFA readout chemistries.

To apply the BINOCULARs, it was also necessary to supply complementary coil peptides each conjugated to ligands for LFA readout. We generated these capture-assisting coil-ligand conjugates by peptide synthesis and click chemistry. Upon activation of toehold switch bispecific reporter expression, a BINOCULAR containing Coil A and Coil B binds to cognate binding partners Coil A* (conjugated to a capture probe, e.g., biotin) and Coil B* (conjugated to a reporter, e.g., FAM), which have been supplied to the reaction (Figure 1). The resulting complex is applied to a lateral flow strip together with ChonBlock buffer, where captured reporters accumulate at the test line to generate a visible signal. Specifically, we first used the fusion Coil P1 - GS linker - Coil P4S (BINOCULAR-P1-P4S) as output for the mycoplasma sensors, while Coil P10 - GS linker - Coil P12 (BINOCULAR-P10-P12) was used as output for the PUM2 sensors. To interface with lateral flow, we conjugated to biotin the cognate coils P2 and P9, which bind to P1 and P10, respectively, which enables the complex to bind to the streptavidin embedded on the test line. Similarly, cognate coils P3S and P11, which bind to P4S and P12, respectively, were conjugated to 6-carboxyfluorescein (FAM) to capture anti-FAM-gold nanoparticles for readout (Figure 2d). A final concentration of 100 nM of each ligand-conjugated coil proved to be the optimal concentration for the biotin conjugating complexes and was used in the experiments.

To ensure a direct comparison of performance, the BINOCULARs were integrated with the same toehold switch sensors previously utilized for the LacZ and LacZα output evaluations (Figures 2b and 2c), replacing the enzymatic outputs. We found that BINOCULARs provided 10- to 31-fold visible signal change after only a 5-minute cell-free reaction (Figure 2d). ImageJ was used to quantify the test band intensities on lateral flow strips, while a strong band can be observed on lateral flow strips by the naked eye (Figure 2e). Additionally, by leveraging the established modularity of coiled-coil libraries, this approach allows sensing elements and functional moieties to be combined interchangeably, which is a key design feature for facilitating integration into versatile downstream applications.

### Integration of LAMP Amplification for High-Sensitivity Detection

Even though chronic mycoplasma contamination is present at high titers (107–108 CFU/mL) in non-turbid cultures, the inability to visibly distinguish the presence of mycoplasma makes earlier detection challenging. Therefore, we designed our platform to meet the stricter regulatory limit of detection (LOD) of ≤ 100 genomic copies/mL,^26^ benchmarking it against gold-standard testing methods. To achieve this level of sensitivity, we incorporated Loop-Mediated Isothermal Amplification (LAMP) combined with a rapid heat-extraction protocol to pre-amplify targets directly from cell culture media.

We screened several extraction methods and amplification primer sets to determine the optimal workflow (Supplementary Figure 2). The best-performing method was selected for integration with the cell-free detection assay (Figure 3a). Cell culture media supernatant was directly taken from the flask or dish and heated up to 100°C for 5 minutes for DNA extraction. The resulting lysate was then directly added to the LAMP reaction and incubated at 61°C for 20 minutes.

**Figure 3.**
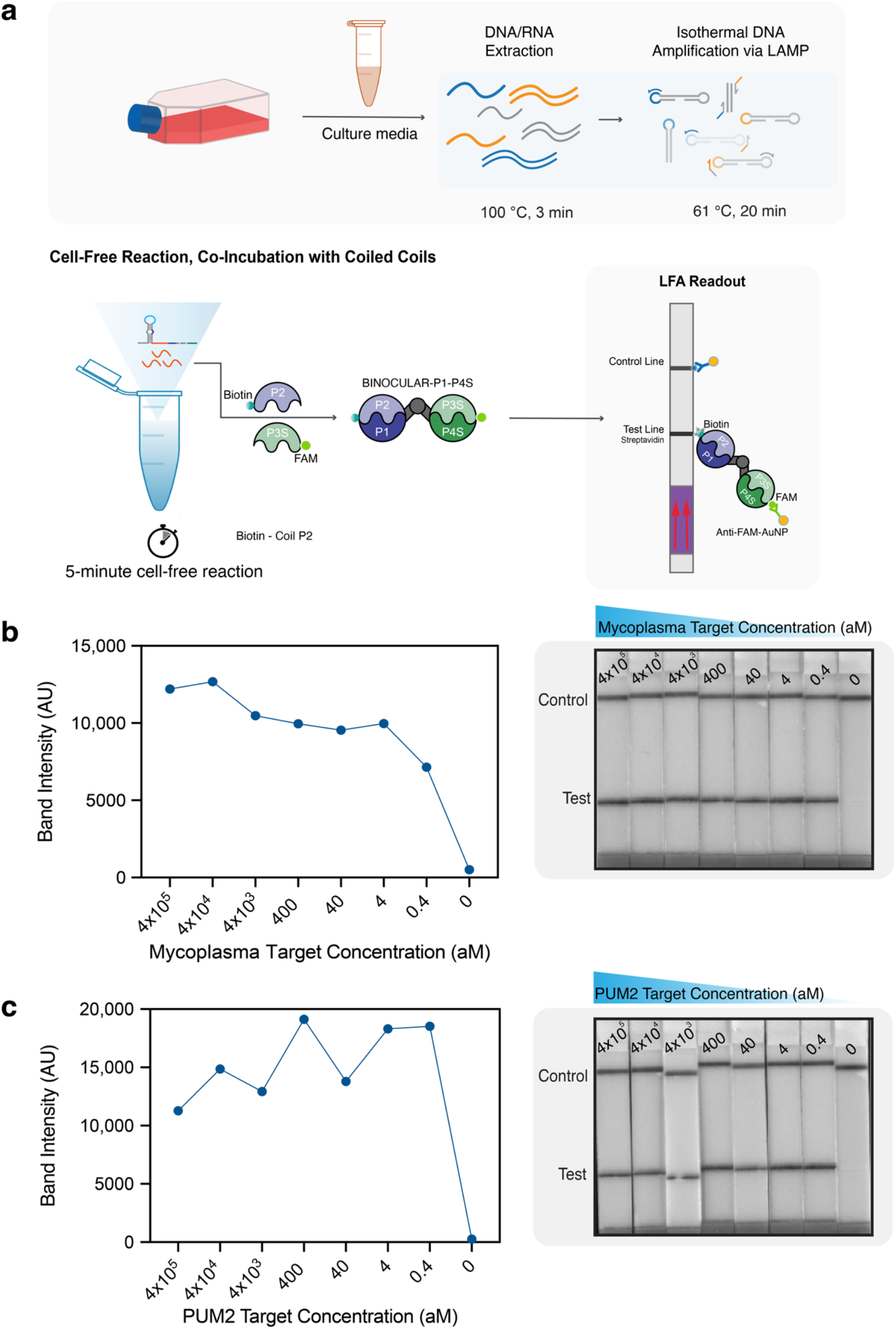
Detection of mycoplasma and control target by coupling with LAMP. (a) Schematic of the integrated mycoplasma detection workflow. Mycoplasma and PUM2 control targets are extracted from cell culture media (taken directly from the flask) by a brief heat-extraction step at 100°C for 3 min. The extracted sample is then diluted 1:1000 in nuclease-free water and added to a LAMP reaction for a 20-minute amplification. 1 µL of the LAMP product is applied to the toehold switch-BINOCULAR detection assay for a 5-minute incubation, followed by lateral flow readout. (b, c) Performance of a representative mycoplasma sensor (b) and a representative PUM2 sensor (c) against serially titrated concentrations of cognate synthetic target. The assay demonstrates a limit of detection (LOD) of 0.4 aM (∼1 copy/µL). The left panel presents the quantified band intensity, while the right panel displays the corresponding lateral flow strips for qualitative visualization.

For LoD determination, we amplified serial dilutions of synthetic target DNA ranging from 400 fM down to 0.4 aM. The resulting amplicons were transferred directly into the cell-free reaction. All six top-performing sensors for both the *Mycoplasma 16S rRNA* and the *PUM2* internal control successfully detected concentrations as low as 0.4 aM (∼1 copy, corresponding to 50 copies/mL). While the system is designed to detect the DNA version of *Mycoplasma 16S rRNA*, when using a LAMP/RT-LAMP kit, the reaction is also able to simultaneously amplify the higher abundant RNA pool of *Mycoplasma 16S rRNA*, of which there are thousands of copies per cell compared to only 1–2 genomic copies^38,39^. This ensures that the assay provides a functional sensitivity significantly higher than the regulatory threshold of 100 genomic copies/mL. Strong visual signals were observed on lateral flow strips for both the *Mycoplasma 16S rRNA* sensor (Figure 3b, Supplementary Figure 3a) and the *PUM2* sensor (Figure 3c, Supplementary Figure 3b) within 5 minutes of applying the mixture of the cell-free reaction product, ligand-functionalized coils, and ChonBlock buffer.

### Multiplex Detection via Modular Coiled-Coil Reporters

We next sought to leverage the modularity and orthogonality of the BINOCULAR system to enable multiplexed nucleic acid detection. In addition to the mycoplasma and internal control targets, we adapted a SARS-CoV-2 sensor previously developed by our group to incorporate a coiled-coil output, demonstrating simultaneous three-plex detection^40^.

To achieve signal discrimination, we cloned distinct pairs of coiled-coils for each BINOCULAR output. We then functionalized the cognate binding partners as follows: the cognate coil peptide for the reporter’s N-terminus was conjugated with a capture ligand (Biotin, 6-TAMRA, or DIG), while the cognate coil peptide for the reporter’s C-terminus was modified with a universal reporter ligand (FAM) (Figure 4a). We investigated various coiled-coil combinations and selected the optimal coil-ligand pairings for the final assay (Supplementary Figure 4). Specifically, the mycoplasma channel utilized the BINOCULAR-P1-P4S with the biotin ligand; the internal control channel utilized the BINOCULAR-P10-P12 coil pairs with the 6-TAMRA ligand; and the SARS-CoV-2 channel utilized the BINOCULAR-P6-P8 with the DIG ligand.

**Figure 4.**
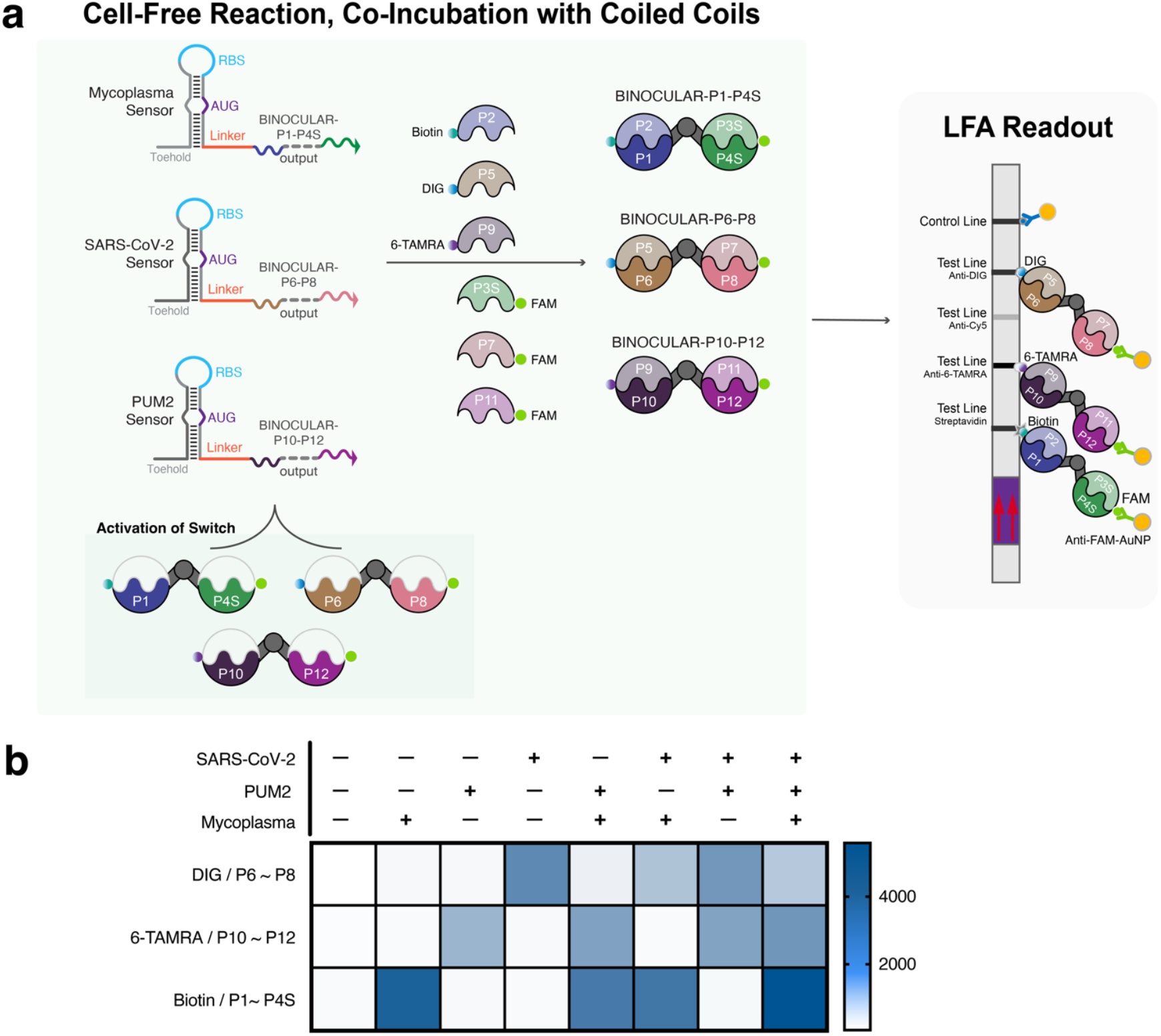
Multiplexed detection of Mycoplasma, PUM2, and SARS-CoV-2. (a) Schematic illustration of multiplexed assay components for three-channel lateral flow readout. Toehold switches for mycoplasma, PUM2, and SARS-CoV-2 were coupled with three orthogonal BINOCULAR outputs and six accompanying coil-ligand conjugates. (b) Heat map of band intensities for detection of three targets in a one-pot cell-free reaction. Coiled-coil pairs P1-P2 and P3S-P4S were used for mycoplasma sensors; P5-P6 and P7-P8 were used for SARS-CoV2 sensors; P9-P10 and P11-P12 were used for PUM2 sensors. Final concentration of the target is 5 µM for all experiment groups. Results are representative of *n = 3* independent technical replicates.

For simultaneous three-plex detection, a master mix containing all three sensors was added to the cell-free reaction and incubated with various target combinations at 37°C. While shorter incubations yielded detectable signals, 15 minutes was selected to maximize signal intensity without significantly prolonging the assay. Subsequently, the ligand-functionalized capture coils were added to the reaction at optimized final concentrations (100 nM for biotin pairs; 250 nM for 6-TAMRA pairs; 200 nM for DIG pairs) prior to the insertion of the lateral flow dipsticks.

We demonstrated that in the presence of a specific target, only the corresponding sensor was activated, generating a detectable band at the appropriate test line (Figure 4b). Furthermore, when multiple targets were present simultaneously, all corresponding sensors responded efficiently, generating distinct bands for a clear multiplexed readout.

### Validation in Complex Biological Matrices and Shelf-Life Stability

To verify the robustness of the platform for practical biomanufacturing and laboratory scenarios, we challenged the assay with various types of contaminated mammalian cell lines and different culture matrices. Several sets of human iPSCs (in StemMACS), HEK 293T cells (in complete Dulbecco’s Modified Eagle Medium), SK-OV-3 cells (in McCoy’s 5A medium), CHO cells (in EX-CELL CD CHO Fusion medium), and synthetic targets (in IDTE buffer pH 8.0) in their corresponding matrices were tested (Figure 5a). Following the standard workflow (Figure 3a), our assay successfully detected the presence of both mycoplasma and the PUM2 control target across all contaminated cultures, regardless of the contamination level.

**Figure 5.**
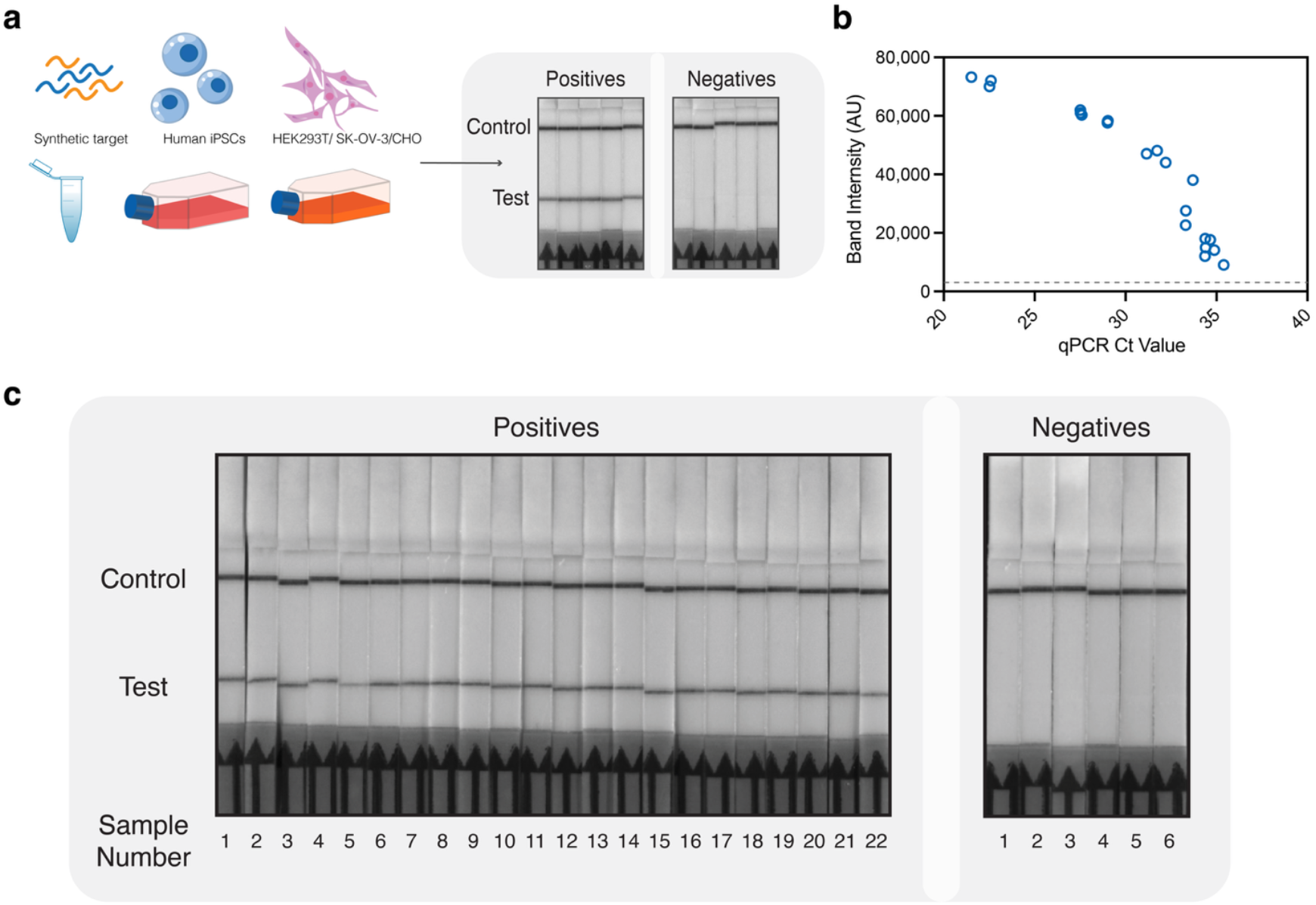
Assay performance validation: matrix compatibility, qPCR correlation, and freeze-dried stability. (a) Matrix and cell compatibility of the toehold switch assay. Detection of mycoplasma in various contaminated cell lines cultured in different media. The lateral flow strips (from left to right) show representative results for a synthetic target in IDTE buffer, human iPSCs in in StemMACS medium, HEK293T cells in complete Dulbecco’s Modified Eagle Medium (DMEM), SK-OV-3 cells in McCoy’s 5A medium, and CHO cells in EX-CELL® CD CHO Fusion medium. (b) Correlation between qPCR Ct values and LFA band intensities. Results are shown for a diverse set of mycoplasma-contaminated cell culture media samples. The analysis demonstrates a strong positive correlation, with a Pearson coefficient (r) of 0.973 (p < 0.0001). The horizontal dashed line indicates a band intensity visible by eye. (c) Performance of the freeze-dried assay for on-site use. Representative LFA results show robust mycoplasma sensor performance 35 days post-freeze-drying. The assay was tested against 1 µL of LAMP product amplified from a 1 copy/µL target. In a test panel of 22 PCR-verified positive samples and 6 PCR-verified negative samples, the freeze-dried assay demonstrated 100% sensitivity and 100% specificity.

We further benchmarked the performance of the mycoplasma sensor on these biological samples against standard qPCR (Figure 5b). The results demonstrated strong agreement, yielding a Pearson correlation coefficient of 0.973 (*p* < 0.0001), while significantly reducing assay time and sample handling requirements.

Beyond static benchmarking, we demonstrated the utility of the assay for the early detection and longitudinal monitoring of contamination in iPSCs cultures. The platform successfully identified mycoplasma as early as the second day post-infection and consistently detected the presence of mycoplasma over the following days (Supplementary Figure 5).

To demonstrate the feasibility of convenient on-site application, we lyophilized the assay components and left them at room temperature with silica gel desiccation packages, avoiding light. After 35 days of storage, the stability of both the mycoplasma (Figure 5c) and PUM2 (Supplementary Figure 6) sensors we evaluated. The lyophilized pellets were reconstituted with 9 µL of nuclease-free water and 1 µL of LAMP amplicons of 1 copy/µL target. The assay maintained its original limit of detection with 10 minutes of reaction time, with 100% sensitivity and 100% specificity across 22 positive samples and 6 negative samples.

Compared to qPCR, the total assay duration was condensed from several hours to approximately 25 minutes. While this platform eliminated the requirement for specialized qPCR instrumentation, the shelf-stable, lyophilized format also removes the need for cold-chain storage. By requiring only reconstitute the lyophilized components with amplicons and functionalized coiled-coil peptides, the procedure minimizes small-volume pipetting iterations, then further mitigates the risk of operator-induced cross-contamination and technical inconsistency.

## DISCUSSION

In this study, we introduced a biosensing platform that employs a bispecific protein reporter for lateral flow readout, demonstrating its capacity for rapid, highly specific, and multiplexed isothermal detection of nucleic acids. The BINOCULAR design, incorporating compact coiled-coil peptides, provides a remarkably fast readout time in as little as 5 minutes of cell-free expression. We found that this readout time is ≥12-fold faster than comparable reactions using standard cell-free reporters like fluorescent proteins or LacZ^6^, and ≥6-fold faster than split lacZ output^7^ while generating negligible non-specific background signal.

Through integration with isothermal amplification, the assay enables the visual detection of the adventitious agent mycoplasma nucleic acids at abundances as low as 1 copy from varying sample matrices. This performance enables early contamination detection and could prevent financial losses and product quality crises in biomanufacturing applications. Furthermore, by demonstrating the assay’s compatibility with lyophilization and its robust performance following long-term ambient storage, we have established its viability as a convenient, cost-effective platform for routine, on-site contamination testing.

Previous translation-based sensors have also faced a trade-off between minimizing equipment requirements and maximizing multiplexing capacity. For instance, enzymatic colorimetric readouts are inherently restricted by the distinguishable colors that enzyme and substrate combinations can produce^14^. However, by leveraging libraries of orthogonal coiled coils, BINOCULAR provides significant improvements in multiplexing. We demonstrated the simultaneous, 3-plexed detection of nucleic acid targets, proving that multiple targets can be identified in a single test without complex instrumentation. We expect that the number of readout channels can be expanded by using additional orthogonal coils and LFA capture chemistries.

In conclusion, our BINOCULAR based system overcomes major limitations of previous translation-based sensors. It achieves a unique synergy of an ultrafast time-to-result, robust multiplexing, and exceptional sensitivity. This performance is enabled by integrating the specificity and programmability of toehold switches, the modularity of coiled-coil peptides, and the portability of LFA readout systems. We propose that this combination, along with its proven cell-free compatibility and operational robustness, establishes it as a powerful platform for adventitious agent detection in biomanufacturing and for a broader range of pathogen diagnostics.

In the future, we expect that BINOCULAR’s multiplexing capacity can be expanded using orthogonal coiled-coil libraries generated by *de novo* design^41^ and by engineering trimeric coiled-coil reporters^42,43^. In addition, the range of analytes detectable using BINOCULAR cell-free reactions could be extended to non-nucleic acid analytes^17^, such as small molecules. These developments will help broaden the utility of BINOCULARs, unlocking their potential for diverse diagnostic and research applications.

## MATERIAL AND METHODS

DNA oligos were purchased from Integrated DNA Technologies (IDT) or Twist Bioscience. Q5 High-Fidelity 2X Master Mix, Gibson Assembly® Master Mix, PURExpress® In Vitro Protein Synthesis Kit, DpnI restriction enzyme and WarmStart® LAMP Kit (DNA & RNA) were purchased from New England Biolabs (NEB). Juice− Cell-free kit was purchased from Liberum Bio. HybriDetect Universal Lateral Flow Assay Kits were purchased from Milenia Biotec. Protector RNase Inhibitor was purchased from Sigma Aldrich.

### Designing and Screening Toehold Switch Sensors

A NUPACK-based selection algorithm described previously^4,32,33^ was used to identify toehold switches for detection of *Mycoplasma* 16S rRNA and PUM2 control. We obtained 48 candidate toehold switches for both targets and performed screening with LacZ output. Top-performing sensors were selected based on ON/OFF ratio after 2 hours of cell-free reaction.

### Toehold Switch Plasmid Construction

The list of plasmids used in this work are provided in Supplementary Table 2. For the sensor screening process, candidate toehold switch sequences were purchased as ultramer DNA strands from IDT, amplified by PCR with Q5 2x Master Mix (New England Biolabs M0492L) then cloned into a pColA backbone (Addgene plasmid #11882) with LacZ output by Gibson assembly.

For testing LacZα output sensors, a sequence encoding the LacZα peptide was obtained by PCR from Addgene plasmid #118819 and cloned into the top-performing sensor plasmids via Gibson assembly to replace *LacZ*.

Gibson assembly used unpurified insert switches or target PCR products and a custom 2x enzyme mix containing 4 U/µL T5 Exonuclease (New England Biolabs M0663S), 25 U/µL Phusion polymerase (New England Biolabs M0530S), 4 U/µL Taq ligase (New England Biolabs M0208L), and 7 ng/µL ET SSB (New England Biolabs M2401S) in Gibson buffer [1 mM dNTPs (New England Biolabs N0446S), 5 mM NAD+ (New England Biolabs B9007S), 500 mM Tris-HCl (VWR 101641-844), 50 mM MgCl2 (Sigma 63069), 50 mM DTT (Sigma 43816), 25% PEG-8000 (Sigma P2139)].

The cloning of sensors with BINOCULAR output was based on the LacZ output sensors. Coiled-coil peptide sequences were obtained from literature^25^ and linked via a flexible glycine-serine (GS) linker. The divalent output sequences were synthesized by Twist Biosciences as double-stranded DNA and then cloned into the top-performing sensor plasmids via Gibson assembly to replace *LacZ*. The following BINOCULAR plasmids will be available from Addgene: pColA-Mycoplasma-TS-BINOCULAR-P1-P4S (Addgene, 257830), pColA-PUM2-TS-BINOCULAR-P10 -P12 (Addgene, 257831), pColA-SARS-CoV-2-TS-BINOCULAR-P6-P8 (Addgene, 257832). Sequences of the BINOCULAR genes are provided in Supplementary Table 3.

Gibson assembly products were transformed into chemically competent *E. coli* DH5α, followed by a 30 min recovery at 37°C in SOC media (New England Biolabs B9020S) shaking at 250 rpm. Cells were plated on LB agar (Sigma L3147) with appropriate antibiotics and grown overnight at 37°C. Plasmids were purified with QIAprep Spin Miniprep Kit (27104). All plasmids were verified by Sanger sequencing.

### Conjugation of Synthetic Coil Peptides

Single coiled-coil peptides were synthesized with an azido group on either the N- or C terminal by AbClonal (Supplementary Table 4). For readout on lateral flow strips, coils were conjugated either with FAM, to enable capture of anti-FAM-labelled AuNPs, or with ligands to promote capture at LFA-embedded test lines. Different DBCO containing ligands (BroadPharm and Integrated DNA Technologies) were bioconjugated to the coiled coils via copper-free click chemistry as described in Supplementary Table 5. The azido-coil peptide and DBCO-containing ligands were incubated at equal molar ratios in 1x PBS buffer overnight at 4°C.

To confirm successful click chemistry, DBCO depletion was monitored by absorbance at 310 nm before and after incubation. The resulting conjugated coils were stored at −20 °C.

### Isothermal Amplification of Targets

LAMP primers were designed using NEB LAMP primer design tool (version 1.5.1) for the template sequence in Supplementary Table 1b. LAMP reactions were prepared according to the manufacturer’s instructions (NEB, E1700L) in a total volume of 20 µL using a full set of 6 LAMP primers (FIP and BIP: 1.6 µM each, F3 and B3: 0.2 µM each, LF and LB: 0.4 µM each) and using 8 µL of sample. Reactions were incubated at 61°C for 20 minutes, followed by 80°C for 5 minutes to deactivate the enzyme. The best primers were selected by verifying the amplicon band intensity on 1.5% agarose gel.

### Toehold Switch Assay with LacZ Output

Sensor screening was performed in cell-free reaction using New England Biolabs PURExpress In Vitro Protein Synthesis Kit (E6800L). Cell-free reactions were prepared with the following components by volume: cell-free solution A, 40%; cell-free solution B, 30%; RNase Inhibitor (Roche, 03335402001, distributed by MilliporeSigma), 2%; chlorophenol red-b-D-galactopyranoside (CPRG) (Roche, 10884308001, distributed by MilliporeSigma, 24mg/ml), 2.5%; with the remaining volume reserved for toehold switch DNA, water and 5 µM of synthetic target.

All sensor plasmids were prepared at a 100 ng/µL concentration after purification. Synthetic target was order from IDT as a DNA ultramer. All candidates were screened at a final concentration of 12 ng/µL in a 5 µL cell-free reaction. Each mycoplasma sensor was tested with mycoplasma synthetic target (ON state) and with a PUM2 gene synthetic target (OFF state), and vice versa for the PUM2 sensors. A BioTek Synergy Neo2 multimode microplate reader (Agilent) pre-heated to 37°C was used for monitoring the cell-free reaction progress for 5 hours.

The signal fold change was calculated by dividing the relative OD 570 of ON state by OFF state. The signal fold change was measured at 2 hours of cell-free reaction for assessment of the toehold switches. Sensors for each target that have >2 ON/OFF were selected and verified in triplicate reactions. Signal at 2 hours was taken for fold change and signal difference significance calculation. The top 3 best-performing sensors from the triplicate tests were selected for future experiments and optimization.

For LacZα output sensors, the lacZω peptide was added to the reaction at a final concentration of 2 µM while the rest of the component concentrations remained the same.

### Toehold Switch Assay with BINOCULAR Output

For single target detection assay, a 10-µL cell-free assay was prepared as a conventional toehold switch assay with either synthetic target or LAMP amplicon. The cell-free reaction was incubated at 37°C for a desired amount of time. For multiplexed detection, a 10-µL cell-free assay was prepared as a conventional toehold switch assay with either synthetic targets or LAMP amplicons. The sensor plasmids concentrations were unified to 200 ng/µL to reserve more volume for multiple sensors and targets. Same final concentration was used for all sensors.

To examine the signal on lateral flow, the cell-free product with BINOCULARs was mixed with ligand-conjugated coils then ChonBlock Blocking/Sample Dilution ELISA Buffer (Chondrex) was added up to a final volume of 50 µL. The corresponding coils were added to the reaction to the final concentration as described in results. The lateral flow strip (Milenia Biotec) was added to the PCR tube and allowed to develop for 5 minutes. Lateral flow strips were imaged using a GelDoc Go Gel Imaging System (BioRad) using colorimetric mode and an exposure time of 0.15 seconds.

### Assay Lyophilization and Preservation

The cell-free assay containing PURExpress reagent and sensor was prepared by either flash freezing in liquid nitrogen or freezing at -80°C for 2 hours, followed by ≥17 hours of freeze-drying at -50°C and 0.04 mbar (Labconco FreeZone Triad Benchtop Freeze Dryer). The freeze dryer was pre-cooled to -50°C or colder before preparing the cell-free reaction mix.

The freeze-dried assay was preserved in sealed ziplock bags with silica gel desiccant packets at room temperature for long-term storage. The freeze-dried assay was reconstituted with 10 µL nuclease-free water and then subjected to the same toehold switch assay reaction steps described above.

### Cell Culture

Human induced pluripotent stem cells (hiPSCs) were maintained in mTeSR Plus medium (STEMCELL Technologies) and cultured on Cultrex-coated plates under standard incubator conditions (37°C in 5% CO2). Cells were passaged every 3-4 days using Accutase. For thawing and routine passaging, mTeSR Plus was supplemented with 5 µM ROCK inhibitor. For cell pelleting, cells were dissociated with Accutase and centrifuged at 200g for 5 minutes. Supernatants were carefully removed, and cell pellets were processed immediately or stored until further use at -80°C.

Human embryonic kidney (HEK293T) cells were maintained in Dulbecco’s Modified Eagle Mediu (Gibco™, 11995065) supplemented with 10% fetal bovine serum (FBS) and 1% Penicillin-Streptomycin (Pen-Strep). HEK293T cells were cultured on tissue culture-treated plates and incubated at standard incubator conditions (37°C in 5% CO2). Cells were passaged every 3 days using 0.25% Trypsin-EDTA (Gibco™, 25200114) according to standard passaging procedures.

### Mycoplasma Contamination

To model mycoplasma exposure, hiPSC culture media was collected to ensure the cells were negative for mycoplasma. Once media was collected, hiPSCs were intentionally contaminated with mycoplasma-positive conditioned media at varying ratios, 24 hours after passaging. After 48 hours of incubation in mycoplasma-positive media supplemented with mTeSR, the mycoplasma-positive media was removed and replaced with fresh mTeSR Plus. Culture media was collected daily for seven consecutive days. Mycoplasma positivity was assessed using the Lonza MycoAlert® Mycoplasma Detection Kit, following the manufacturer’s instructions as a reference. On day 7 post-contamination, cells were pelleted for downstream analyses.

### qPCR Determination of Mycoplasma Copy Numbers

A standard curve was established with ssDNA purchased from IDT. Various copy numbers were diluted from a stock of 100 µM. Each 20µL reaction contained 10 µL Luna Universal Probe qPCR Master Mix, forward primer (10 µM), reverse primer (10 µM), FAM probe (10 µM), various amount of template and Nuclease-free Water. A StepOnePlus Real-Time PCR instrument was used for the qPCR experiment with the following settings: initial denaturation at 95°C 1 minutes, 40 cycles denaturation at 95°C for 15 seconds, anneal/extend/read at 60°C for 30 seconds. Ct values of each sample were recorded for comparison with the cell-free assay.

### Statistical Analysis

ImageJ was used to quantify the test band intensities on lateral flow strips. All OD 570 and band intensity data were graphed and interpolated in GraphPad Prism. GraphPad Prism was also used to plot mean values and calculate the correlation statistics.

## Author Contributions

Y.L., X.W., D.A.B., and A.A.G. conceived the study. Y.L. and N.R.A. developed the methodology, designed, performed the experiments and data analysis. A.E. and S.N. performed the experiments. Y.L. and C.F. acquired the cell culture medium samples. N.R.K. provided sequences and contributed to the optimization of LFA experiments. A.A.G., D.A.B., and X.W. supervised the research and acquired funding. Y.L. and A.A.G. wrote the manuscript.

## Acknowledgements

This research was sponsored by the Office of the Secretary of Defense and was accomplished under Agreement Number W911NF-17-3-0003. This work was also supported by Defense Advanced Research Projects Agency (DARPA) funding (Contract No. N66001-23-2-4042). N.R.A. was supported by the NIH Training Program in Synthetic Biology and Biotechnology (1T32GM130546). The views, opinions, conclusions, and/or findings contained in this document are those of the authors and should not be interpreted as representing the official views or policies, either expressed or implied, of the Department of Defense, the Office of the Secretary of Defense, or the U.S. Government. The content is solely the responsibility of the authors and does not necessarily represent the official views of the National Institutes of Health. The U.S. Government is authorized to reproduce and distribute reprints for Government purposes notwithstanding any copyright notation herein.

