## Supplementary Figures for "Bispecific Coiled-Coil Reporters Enable Rapid, Multiplexed Detection of Mycoplasma via Toehold Switch Sensors"

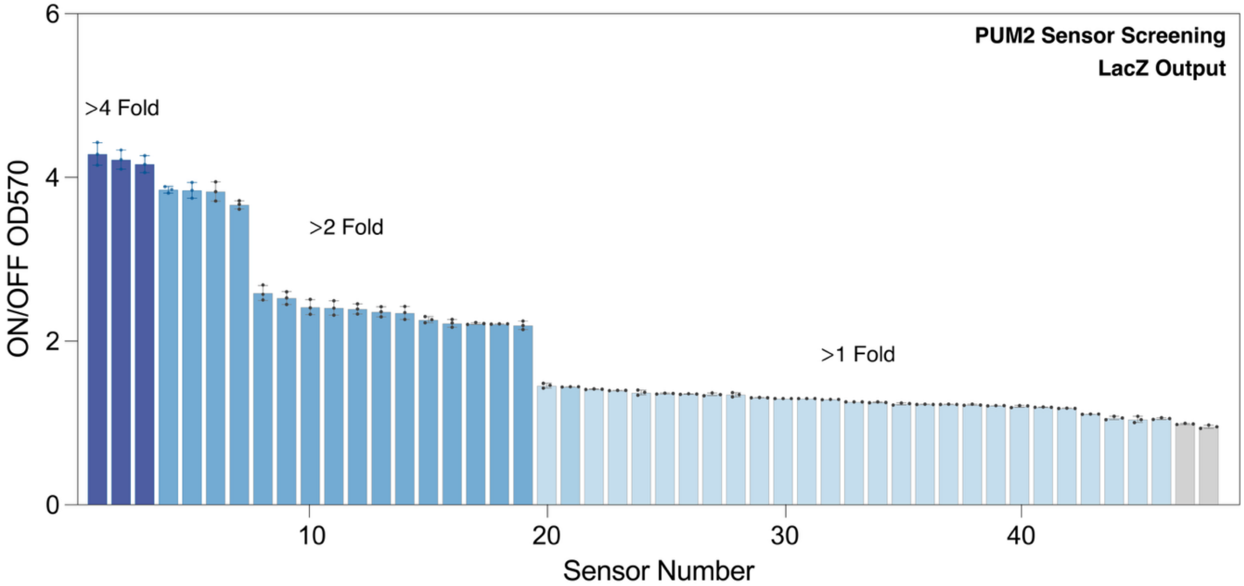


**Supplementary Figure 1.** Screening of PUM2 toehold switch sensors with LacZ output. OD570 fold change obtained 2-hour of cell-free reaction for 48 toehold switch sensors determined in the presence or absence of 5 µM of cognate *PUM2* target.

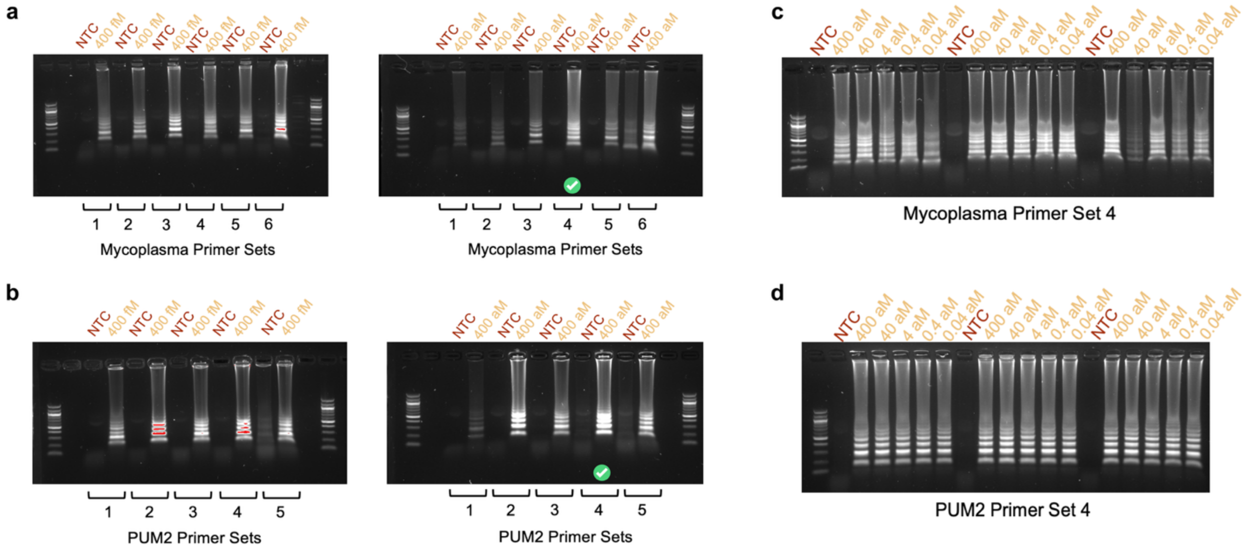


**Supplementary Figure 2.** LAMP primer characterization. Gel images show the initial primer screening results for both *Mycoplasma 16S rRNA* (a) and *PUM2* (b) at 400 fM (high) and 400 aM (low) target concentration. We then examined on the limit of amplification of selected primers for both *Mycoplasma 16S rRNA* (c) and *PUM2* (d) in triplicate reactions.

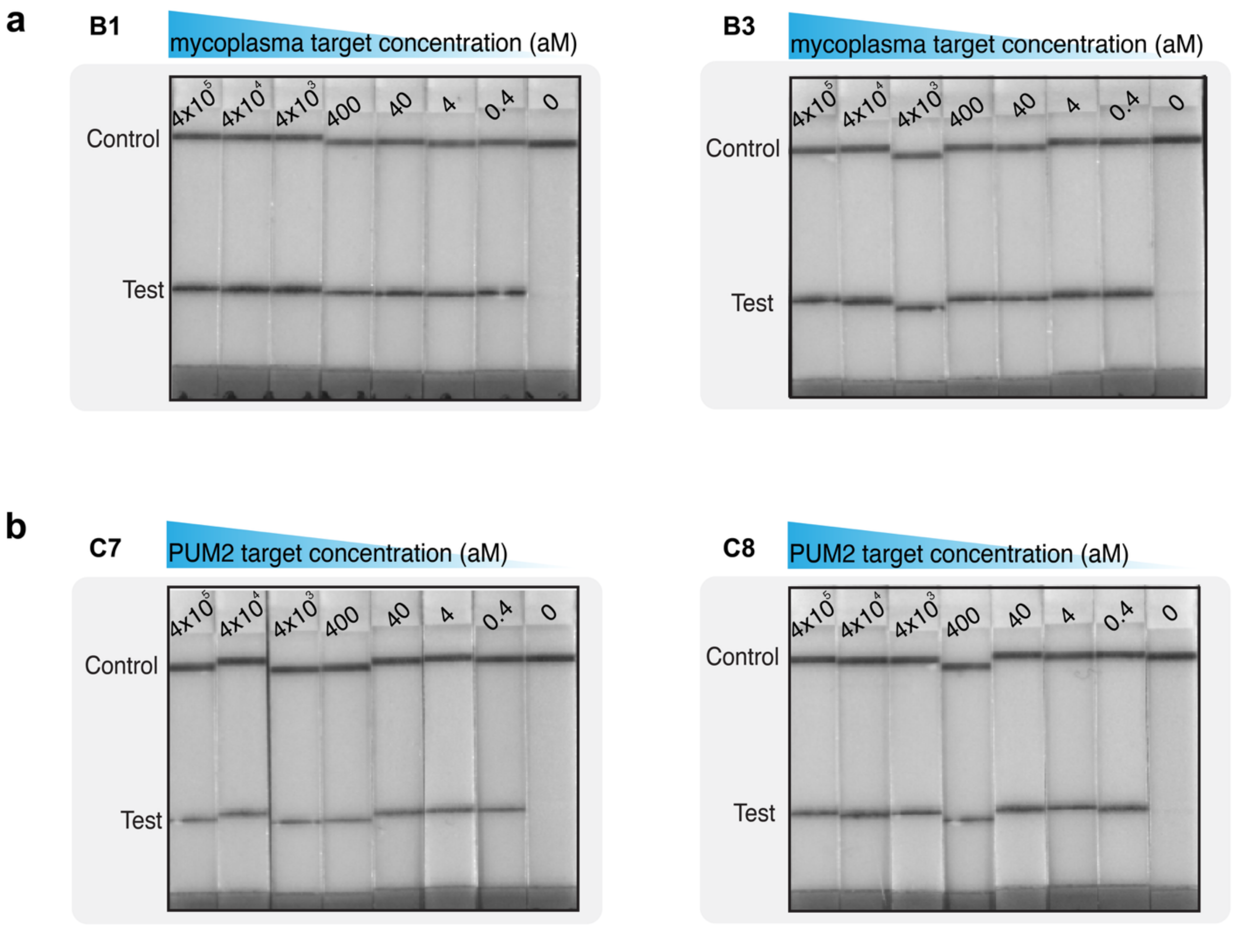


**Supplementary Figure 3.** Performance of two top-performing sensors B1 and B3 for *Mycoplasma 16S rRNA* (a) and sensors C7, C8 for *PUM2* (b) against serially titrated concentrations of cognate target.

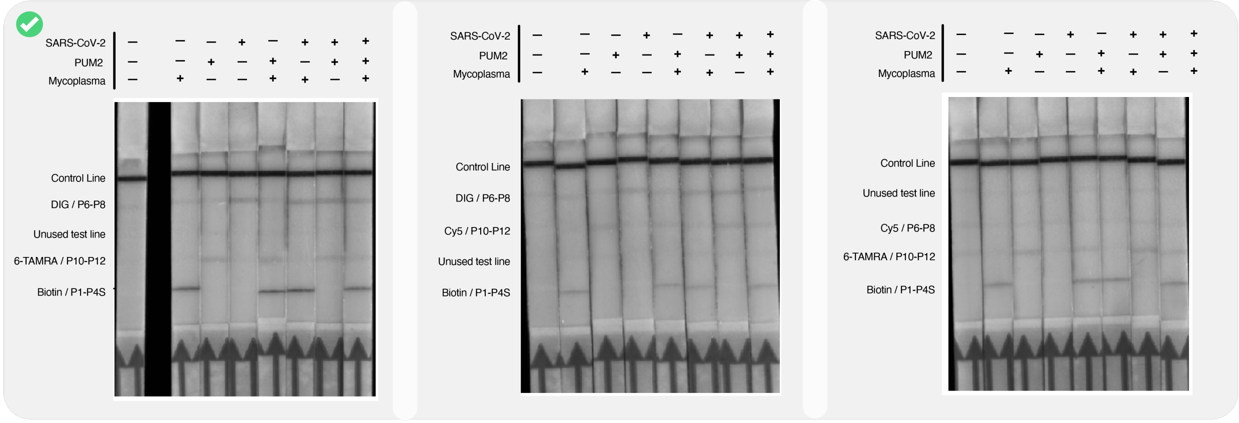


**Supplementary Figure 4.** Various coiled-coil combinations were tested for 3-plex detection, and the optimal coil-ligand pairings, and the best concentration ratios were selected for the final assay. Representative results from three top-performing conditions. For example, the image on the left shows the use of P3S-**Biotin** as the capture conjugate, coupling with FAM-P2 and BINOCULAR-P1-P4S for *Mycoplasma 16S rRNA* detection; P9-**6-TAMRA** as the capture conjugate, coupling with P11-FAM and BINOCULAR-P10-P12 for *PUM2* detection; P5-**DIG** as the capture conjugate, coupling with FAM-P7 and BINOCULAR-P6-P8 for SARS-CoV-2 *N* gene detection. Images in the middle and on the right show other combinations.

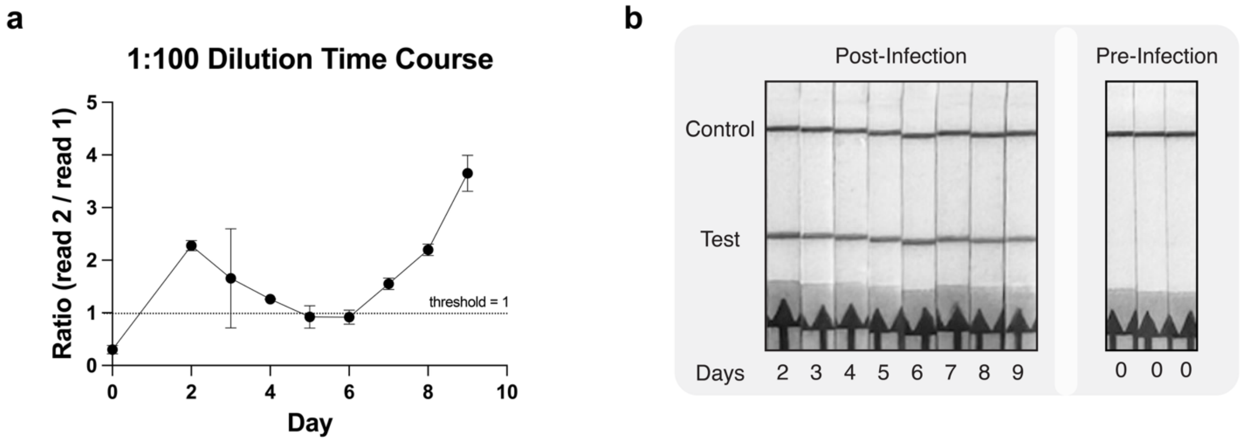


**Supplementary Figure 5.** Early detection and longitudinal monitoring of *Mycoplasma* contamination in hiPSCs cultures. (a) Time-course validation of *Mycoplasma* infection in human induced pluripotent stem cell (hiPSC) media using the Lonza MycoAlert® detection kit. Contamination status is determined by the ratio of Read 2 to Read 1, with a threshold of 1 (dashed line) indicating positivity. Data are shown for 1:100 diluted samples. (b) Representative images of the lateral flow assay strips for samples collected at the same time points. "Pre-Infection" (Day 0) control samples show only the control band, while "Post-Infection" samples (Days 2–9) exhibit a clear test band, confirming the detection of *Mycoplasma* throughout the monitoring period starting from the second day post-infection.

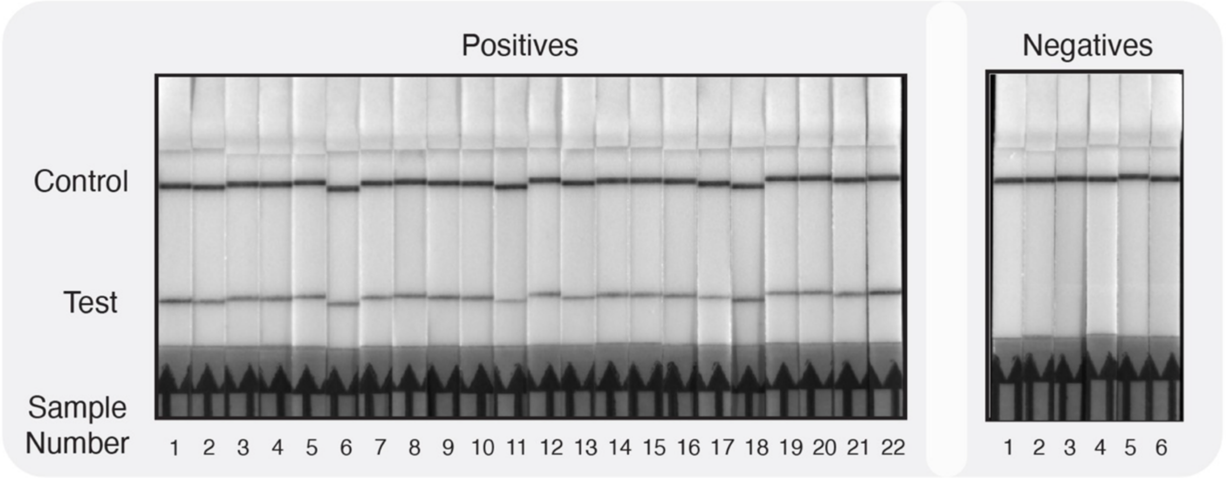


**Supplementary Figure 6.** Performance of the freeze-dried assay for on-site use. Representative LFA results show robust PUM2 sensor performance 35 days post freeze-drying. The assay was tested against 1 μL of LAMP product amplified from a 1 copy/μL target. In a test panel of 22 PCR-verified positive samples and 6 PCR-verified negative samples, the freeze-dried assay demonstrated 100% sensitivity and 100% specificity.
